# Gut Microbiota Signatures of Brain and Peripheral gamma-amino butyric acid (GABA) in Healthy Adults

**DOI:** 10.64898/2026.09.01.748531

**Authors:** Andrea Monteagudo-Mera, Suyi Xie, Valentina Fanti, Claudia Rodriguez-Sobstel, June Christoph Kang, Carolyn B. McNabb, Nicholas Hedger, Jack Newcomb, Nandana Kokkot, Holly Richards, Sasha Dahl, Shan Shen, Anisha Wijeyesekara, Kimon-Andreas Karatzas, Glenn Gibson, Bhismadev Chakrabarti

## Abstract

**Background:** Gamma-aminobutyric acid (GABA), the principal inhibitory neurotransmitter in the mature human nervous system, plays a central role in neurodevelopment, emotion regulation, and sensory processing. Many psychiatric drugs act on the GABAergic system, highlighting its importance for healthy brain function. Individual differences in brain and circulating GABA levels have traditionally been attributed to human genetic variation. However, several gut bacteria produce neuroactive compounds, including GABA, raising the possibility that microbial variation influences peripheral and central GABA levels and related measures in humans. This relationship remains largely unexplored.

**Methods:** In a cross-sectional study of 251 healthy adults, we investigated associations between gut microbiota composition, central and peripheral GABA, and associated features. Central (brain) GABA was quantified using magnetic resonance spectroscopy, peripheral (blood) GABA using enzyme-linked immunosorbent assays, and gut microbiota profiles using absolute and relative abundance measures derived from Flow Cytometry Fluorescent In Situ Hybridisation and 16S rRNA gene sequencing respectively.

**Results:** Motor cortex GABA+ levels were positively associated with greater numbers of *Bacteroides/Prevotella* bacteria. This association was replicated with peripheral GABA. Additionally, peripheral GABA was associated with greater numbers of lactic acid bacteria, *Eubacterium rectale/Clostridium coccoides*, and with predicted microbial gene pathways involved in GABA synthesis. No association was observed between central and peripheral GABA concentrations-suggesting potentially different contributions from microbial metabolism.

**Conclusions:** These findings provide new evidence pointing to factors other than human genetic variation underlying individual variability in central and peripheral GABA. They also set the stage for longitudinal and intervention studies to investigate the mechanisms through which microbiota can contribute to healthy brain function.

## Introduction

The gut microbiota is a key component of the gut-brain axis, influencing host physiology far beyond the gastrointestinal tract. In addition to established roles in immune regulation and metabolic homeostasis, these microbial communities may communicate bidirectionally with the brain through multiple pathways, including the vagus nerve, enteric nervous system, and immune and endocrine signalling. This communication is mediated in part by microbial metabolites, including short-chain fatty acids (SCFAs) as well as neuroactive compounds such as gamma-aminobutyric acid (GABA), serotonin, dopamine, and glutamate. The ability of gut bacteria to produce and modulate neuroactive compounds has attracted attention as it represents an important biochemical mechanism linking the gut microbiota to central nervous system functionality.

GABA is of particular interest because it is the major inhibitory neurotransmitter in the central nervous system and plays an important role in neurodevelopment, sensory processing, motor control, emotional regulation, and cognitive function (1). Some gut bacteria can produce GABA, including strains of *Bacteroides, Bifidobacterium, Lactobacillus* and *Alistipes* (2), or catabolise it as has been reported for *Acinetobacter* (3). Bacteria synthesise and export GABA mainly through their glutamate decarboxylase (GAD) system, which helps them survive acidic environmental conditions such as those in the intestinal tract. Gut microbial communities differ substantially between individuals (4), and can influence central GABAergic function indirectly through vagal, immune, endocrine, and metabolic signalling pathways (5-7). Several neuropsychiatric and neurological conditions characterised by altered GABAergic signalling have also been associated with changes in microbial taxa or metabolic pathways involved in GABA production and metabolism (8-10). However, much of the mechanistic evidence linking gut microbiota to GABAergic function comes from rodent models. For example, administration of *Lactobacillus rhamnosus* significantly increased brain GABA (11) and altered brain GABA receptor expression in a region-dependent manner (5), while administration of *Lactobacillus reuteri* improved stress-related behavioural abnormalities and serum GABA levels in mice (12). Similarly, restoring SCFA significantly changed serum GABA and brain GABA levels in mice with depressive-like features (13). However, rodents differ widely from humans in their microbiota, brain structure, as well as dietary habits (particularly, coprophagy).

In humans, direct evidence remains limited, particularly in general population samples (14, 15). A faecal microbiota transplantation study suggested that manipulating gut microbiota composition of insulin resistant individuals altered circulating GABA levels (16). However, it remains unclear whether this result generalises to healthy adults, and if variation in resident gut microbiota is associated with peripheral and central GABA. A further limitation of previous work is the predominant reliance on relative abundance measures for quantifying microbiota, which do not reflect absolute bacterial counts. To address this limitation in the current study, we use both Flow-FISH-derived absolute quantification as well as 16S sequencing to provide complementary assessments of microbiota composition.

Proton magnetic resonance spectroscopy (1H-MRS) provides a non-invasive and reliable estimate of the level of GABA in a specific brain region (17), while peripheral levels of GABA can be measured reliably using ELISA (18). MRS-estimated brain levels of GABA have also been shown to correlate strongly with motor network functional connectivity in resting state fMRI (19), as well as tactile sensory processing and motor performance (20, 21).

In this cross-sectional study, we used a biochemically informed, multi-level approach to investigate associations between gut microbiota and GABA-related phenotypes in 251 healthy adults. We measured gut microbiota composition using both Flow-FISH-based absolute quantification and 16S rRNA sequencing, blood GABA concentrations using ELISA, and regional brain GABA+ using 1H-MRS.

We hypothesised that the level of GABA-producing bacterial groups will be positively associated with brain and peripheral GABA. The study and its hypotheses were preregistered on https://aspredicted.org/whfh-sdpn.pdf.

## Methods and Materials

### Study design

This was a single-centre, cross-sectional observational study conducted at the University of Reading, UK, between February 2022 and September 2024. Participants were recruited through advertisements at the University of Reading, including campus posters and leaflets, via social media, and through local GP practices. Of 1,618 individuals screened, 613 were eligible, 265 were enrolled, and 251 attended the study visit and were included in the present analysis.

During study visits, participants provided faecal samples for gut microbiota analysis and blood samples for serum GABA quantification. Participants also underwent magnetic resonance spectroscopy (MRS) to quantify brain GABA levels and completed behavioural tasks and questionnaires related to mental health.

The study was conducted in accordance with the Declaration of Helsinki and received ethical approval from an NHS Research Ethics Committee, REC reference 23/WA/0042. Written informed consent was obtained from all participants prior to participation.

### Participants

Participants were eligible if they were male, right-handed, aged 18–50 years, of White European background, raised in the UK or another European country, and had a body mass index (BMI) between 18.5 and 30 kg/m^2^. Inclusion criteria were chosen to minimise systematic confounds due to gender, ethnicity, and geographical background – all of which can influence on gut microbiota composition (22). Participation was restricted to right-handed individuals to minimise brain signal variability from hemispheric lateralisation (23). Participants were excluded if they had used antibiotics or proton pump inhibitors within the previous 3 months; were current smokers or had smoked regularly within the previous 6 months; consumed more than 14 units of alcohol per week; were currently using psychotropic medication or recreational drugs; were taking antibiotics, probiotics or prebiotic supplements; or had a current diagnosis of a neurological, developmental, psychiatric, or any gastrointestinal condition, including inflammatory bowel disease or irritable bowel syndrome. The target sample size was calculated using the *pwr* package in R. A sample of 251 participants provided >95% power to detect correlations of approximately r = 0.25 at a two-sided α level of 0.05, allowing for approximately 10% missing data.

### Procedure

All data were collected during a single study visit at the University of Reading. Participants provided faecal and blood samples, completed questionnaires and behavioural tasks, and underwent MRS to quantify brain GABA levels. Height and weight were measured during the study visit, and body fat percentage estimated using bioelectrical impedance analysis (Tanita BC-418). Participants completed self-reported questionnaires to assess anxiety (State–Trait Anxiety Inventory, STAI-T and STAI-S, respectively), depressive symptoms (Center for Epidemiologic Studies Depression Scale, CES-D) and autistic traits (Autism-Spectrum Quotient, AQ). Participants completed three computer-based behavioural tasks assessing motor performance, emotion recognition, and tactile sensory function. These tasks were included as exploratory behavioural outcomes because performance in these domains has previously been linked to GABAergic function and/or brain GABA levels (21, 24, 25).

### Magnetic resonance spectroscopy

Brain GABA levels were assessed using MRSData were acquired on a 3T Siemens MRI scanner using a MEGAPRESS sequence for edited detection of GABA (26). It should be noted that the edited GABA+ signal represents local GABA together with macromolecules. The test voxels (3×3×3cm) were positioned in the left sensorimotor cortex, corresponding to the hand knob region, and in the occipital cortex centred on the midline. (Figure 1).

**Figure 1:**
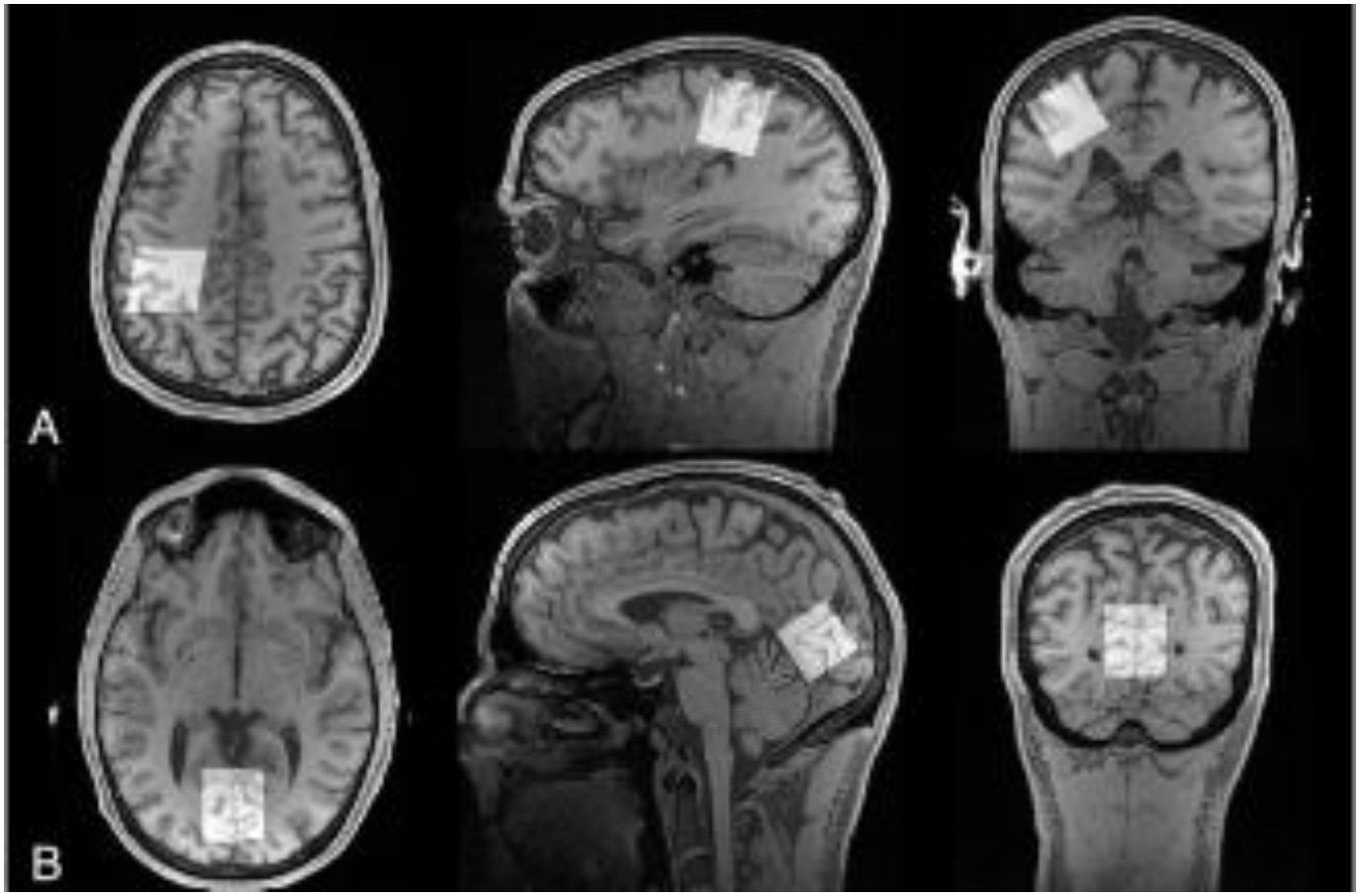
Voxel position in transverse, sagittal and coronal planes in the sensorimotor cortex (A) and occipital cortex (B). MRS data were pre-processed and quantified using Osprey, an open-source MATLAB toolbox for MRS analysis (for analysis scripts see https://github.com/bhismalab/GutBrain) (53). MEGAPRESS spectra were processed using Osprey’s standard pipeline, including frequency and phase correction, eddy-current correction using water reference scans, and linear combination modelling. Spectra were fitted using Osprey’s separate fitting approach with macromolecule modelling enabled using the 1:1 GABA:MM3co model. Because the edited resonance at approximately 3.0 ppm includes contributions from co-edited macromolecules and homocarnosine, GABA+ estimates were extracted and reported. Glx (combined glutamate and glutamine) was also quantified. Metabolite concentrations were quantified using alpha-corrected water-scaled estimates to account for differences in voxel tissue compositions. Further details of spectral processing and modelling are provided in the Supplementary Materials.

### Resting state functional MRI

Resting-state functional MRI (rs-fMRI) was acquired to assess functional connectivity within the sensorimotor network as a proxy measure of GABAergic activity in the motor cortex. Functional connectivity within the motor network was selected as the network of interest because it has previously been shown to be negatively associated with GABA in the motor cortex (19), and to remain stable over time within adults (27). Data were pre-processed with fMRIPrep (version 25.2.3; (28)), followed by computation of functional connectivity using XCP-D (version 0.13.0; (29)). The motor subnetwork was defined using the Schaefer 1056-parcel cortical parcellation (30). Each Pearson correlation in the motor subnetwork (194 × 194 submatrix) was Fisher r-to-z transformed, and the mean was taken across all unique off-diagonal edges to yield a single within-motor-network connectivity value per participant. For more details see Supplementary Materials section 6.

### Peripheral GABA quantification

Serum GABA concentrations were quantified using a commercially available ELISA kit (LDN, Nordhorn, Germany) according to the manufacturer’s instructions. Additional assay details are provided in the Supplementary Materials section 8.

### 16S rRNA gene sequencing and analysis

Approximately 200 mg of faecal material was transferred into PowerBead Pro tubes supplied with the QIAamp PowerFecal Pro DNA Kit (Qiagen, Hilden, Germany). DNA extraction was performed according to the manufacturer’s instructions. Aliquots of extracted DNA were amplified using universal primers targeting the V4–V5 hypervariable region of the bacterial 16S rRNA gene. Sequences were demultiplexed by Novogene Ltd. All samples were initially processed together in QIIME 2 v2025.7 (31) to ensure consistent quality filtering, denoising and ASV generation across the full dataset. Reads were trimmed based on sequence quality, and pre-filtering of sequence contaminants, removal of chimeras, merging of paired-end reads and denoising were conducted using the DADA2 pipeline within QIIME 2. After completing quality filtering steps, 72,892 ASVs were identified across the full dataset of 251 samples. The SILVA 138 99% reference database was used for taxonomic classification of representative ASV sequences. A classifier for the V4–V5 region amplified with the 515F/907R primer pair was trained and used to classify sequences into their respective taxonomic assignments. Diversity analyses were conducted using QIIME 2 to assess microbial community composition and phylogenetic diversity. Further extraction and sequencing details are provided in the Supplementary Materials section 9.

### Flow Cytometry-Fluorescence In Situ Hybridisation (Flow-FISH) for Bacterial Enumerations

Freshly voided faecal samples were diluted in sterile 1x phosphate-buffered saline (PBS; pH 7.3, 0.1M) to prepare 10% (w/v) faecal slurries. Aliquots were centrifuged at 10,000 × g for 10 min, and bacterial pellets fixed in cold 4% (v/v) paraformaldehyde (PFA; Sigma-Aldrich, Poole, UK) for 4–6 h at 4°C. Fixed cells were washed twice with PBS, resuspended in PBS/ethanol, and stored at −20°C until analysis.

Flow-FISH analysis was performed as previously described (32). Total bacteria were quantified using an equimolar mixture of Eub338 I, Eub338 II and Eub338 III probes. Group-specific probes were used to quantify *Bifidobacterium* spp. (BIF), *Bacteroides–Prevotella (*BAC*), Lactobacillus/Enterococcus* (LAB), *Clostridium coccoides–Eubacterium rectale* (EREC), *Roseburia/Eubacterium rectale* (RREC) and *Clostridium histolyticum* groups (CHIS). The absolute cell counts of these groups were log-transformed for analysis.

### Dietary assessment

Dietary intake was assessed using eNutri web-based food frequency questionnaires (FFQ), which estimate habitual food and nutrient intake over the previous 4 weeks (33). Dietary variables were collected to account for potential dietary influences on gut microbiota composition.

### Bioinformatic and statistical analysis

Microbiota analyses were conducted using 16S rRNA gene sequencing-derived relative abundance tables, alongside quantitative bacterial group measurements obtained by Flow-FISH. The 16S rRNA gene was input to PICRUST2 software for metabolic function predictions based on the Kyoto Encyclopedia of Genes and Genomes (KEGG) database (34). The predicted KO enzyme profiles were used to estimate the abundance of 43 Gut-Brain Modules (GBMs), which represent microbial metabolic pathways linked to the production or degradation of neuroactive compounds that may interact with the host nervous system (35). Abundances of genera and GBMs were normalised by centred-log ratio (CLR). Microbiome Multivariable Associations with Linear Models (MaAsLin2) were applied to identify linear associations between taxa, GBMs, and GABA related estimates, controlling for age, body fat percentage, and alcohol intake (36). Spearman correlations were used to assess associations between gut microbiota features, serum GABA, MRS-derived GABA+, as well as behaviour tasks variables. Correlation network was conducted to visualise the interrelationship among them. Sensitivity analysis was conducted by additional adjustment for energy and fibre intake, excluding those with high autistic traits (AQ>32). Benjamini–Hochberg correction method was used to correct the false discovery rate of multiple comparisons. Associations were considered nominally significant at *p* < 0.05 and FDR significant at *q* < 0.25. A *q* <0.25 is the MaAsLin2 default threshold and has been widely used in discovery-phase microbiome studies requiring subsequent validation (35-37). R packages *vegan, maaslin2*, and *igraph* were used for data analysis.

## Results

### Participant characteristics

The cross-sectional sample included 251 participants (see Table 1 for detailed characteristics).

**Table 1.** Participant characteristics.

| **Variable** | **Mean ± SD / n (%)<sup>1</sup>** |
|---|---|
| Age (years) | 34.49 ± 9.18 |
| Body fat (%) | 17.93 ± 5.77 |
| BMI (Kg/m<sup>2</sup>) | 24.38 ± 2.81 |
| Alcohol (g/day) | 13.73 ± 14.78 |
| STAI-S | 32.85 ± 9.65 |
| STAI-T | 40.34 ± 9.95 |
| CES-D | 11.72 ± 8.78 |
| AQ | 19.25 ± 7.36 |

**<sup>1</sup>** Continuous variables are presented in mean and standard deviations. Categorical variables are presented in number and percentage (%). AQ, Autism Spectrum Quotient; CES-D, Center for Epidemiologic Studies Depression Scale; STAI, State-Trait Anxiety Inventory

### Preregistered Analyses

#### Brain and peripheral GABA are associated with partially distinct microbial signatures

Motor cortex GABA+ was positively associated with Flow-FISH quantified *Bacteroides/Prevotella* (adjusted Spearman rho = 0.155, p= 0.018, q = 0.129, Figure 2A, Supplementary Table S1). Motor cortex Glx was negatively associated with *Bacteroides/Prevotella* cell counts (rho = -0.141, p= 0.031, q = 0.111). No significant associations were noted with occipital GABA+.

**Figure 2.**
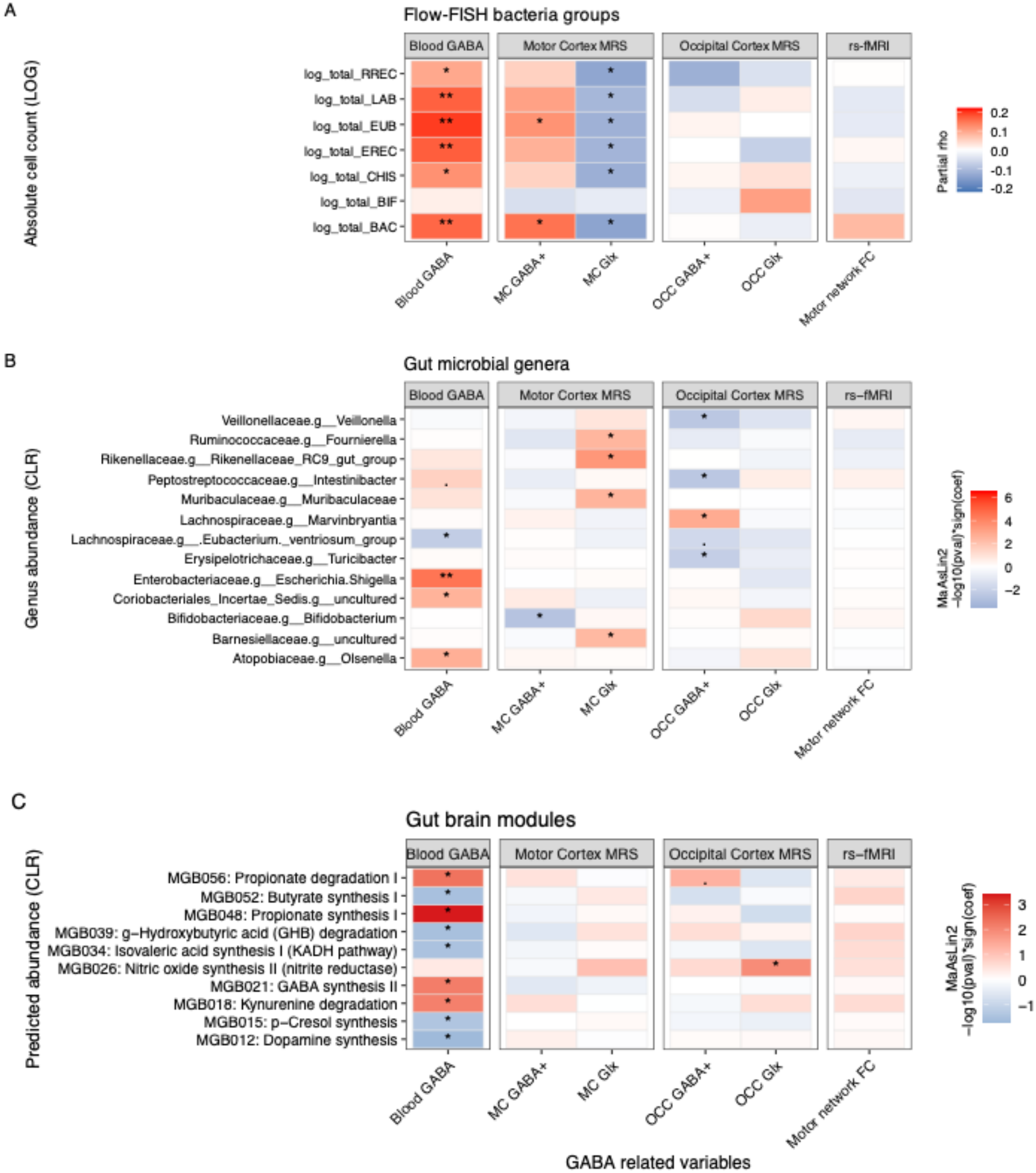
Associations between GABA-related variables and gut microbiota composition, predicted function, and absolute abundance. Heatmaps show associations between peripheral (blood) and brain GABA-related measures and (**A**) absolute cell counts (log-transformed) of Flow-FISH-targeted bacterial groups, derived from partial Spearman correlation analyses; (**B**) abundance (CLR-transformed) of gut microbial genera, derived from MaAsLin2 models; and (**C**) predicted abundance (CLR-transformed) of gut-brain modules (GBMs), derived from MaAsLin2 models. In panel A, colour intensity represents the Spearman correlation coefficient (rho), with red indicating a positive and blue indicating a negative association. Asterisks denote statistical significance based on FDR-corrected q-values (*q < 0.25; **q < 0.05; *** q<0.01), while “·” denotes a nominally significant association (p < 0.05) that did not survive FDR correction.

A similar pattern was noted for peripheral GABA, which was positively correlated with several Flow-FISH quantified groups, including *Bacteroides/Prevotella* (rho = 0.165, *q* = 0.023, *Lactobacillus/Enterococcus* (rho = 0.171, *q* = 0.023), and *Clostridium coccoides–Eubacterium rectale* (rho = 0.173, *q* = 0.023, Figure 2A, Supplementary Table S1).

Abundance of bacterial genera, measured with 16S sequencing provided additional evidence for associations between brain GABA and gut microbial taxa (See Supplementary Table S2). Motor network functional connectivity index was not associated with any measured bacterial genus.

Higher peripheral GABA was positively associated with several 16S derived taxa abundance (Figure 2B; Supplementary Table S2). *Escherichia–Shigella* showed the strongest positive association (*q* = 0.015).

#### Gut microbiota and behavioural tasks related to GABA-ergic processing

No significant association was observed between Flow-FISH and behavioural task performance (Supplementary Table 3). In contrast, several 16S-derived bacterial taxa were significantly associated with behavioural task measures (Figure 3A, Supplementary Table S3). *Sutterela* was negatively associated with reaction time in motor tasks (*q* = 0.038).

**Figure 3.**
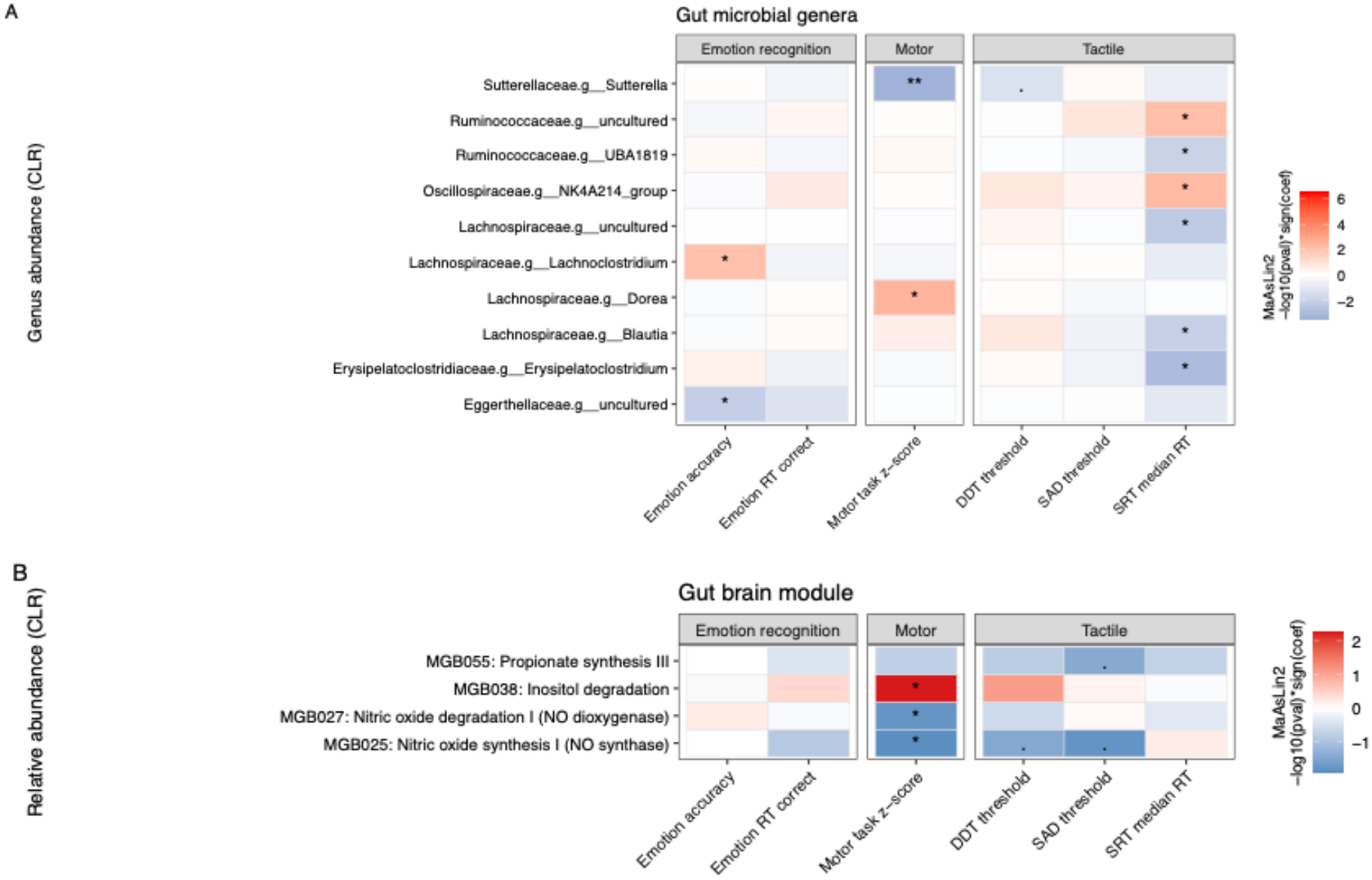
Gut microbial genera and predicted gut-brain modules are associated with performance on behavioural tasks related to GABA-ergic processing. Heatmaps show associations between (**A**) abundance (CLR-transformed) of gut microbial genera and **(B)** predicted abundance (CLR-transformed) of gut-brain modules (GBMs) with behavioural task variables, derived from MaAsLin2 models. Colour intensity represents −log10(p-value) × sign(coefficient), with red indicating a positive association and blue indicating a negative association between microbial/module abundance and the behavioural variable. Behavioural task variables are grouped by domains (Dynamic Detection Threshold [DDT], emotion recognition, motor task, Static Detection Threshold, and Simple Reaction Time [SRT]) along the x-axis. Asterisks denote statistical significance based on FDR-corrected q-values (*q < 0.25; **q < 0.05), while “·” denotes a nominally significant association (p < 0.05) that did not survive FDR correction.

### Exploratory Analyses

#### Microbial Neuroactive Potential Is Associated With Peripheral GABA

Peripheral GABA was positively associated with microbial GABA synthesis pathway (MGB021; q = 0.143). Additionally, it was positively associated with SCFA metabolism modules (MGB048: Propionate synthesis I, *q* = 0.069; MGB056: Propionate degradation I, *q* = 0.143; MGB052: Butyrate synthesis I, *q* = 0.222) (Figure 2C).

MRS-estimated GABA+ levels, or motor network functional connectivity index did not show a significant association with predicted GBM signals after FDR correction (Figure 2C and Figure 3B). Full results are shown in Supplementary Table S5.

#### No relationship Between Peripheral and Brain GABA

Spearman correlation analyses showed that serum GABA was not correlated with motor cortex GABA/GABA+ nor with occipital GABA/GABA+ (Supplementary Table S6).

#### Interrelationship between GABA, gut microbiota and behavioural tasks related to GABA-ergic processing

A correlation network was constructed to visualise associations within one step of blood, brain GABA+ or behaviour outcomes (Spearman |ρ| ≥ 0.15, *p* < 0.05, *q* < 0.10; Figure 4, Supplementary Table S7). Peripheral GABA showed the highest degree of centrality, forming the central hub of the network and correlating with several predicted GBMs (MGB021: GABA synthesis II; MGB056: Propionate degradation I; MGB012: Dopamine synthesis; MGB039: GHB degradation; MGB048: Propionate synthesis I; MGB052: Butyrate synthesis I; MGB018: Kynurenine degradation), genera (*Olsenella, Escherichia–Shigella, Erysipelatoclostridium, Intestinibacter, UBA1819, NK4A214 group*, and uncultured genus within *Coriobacteriales*), and bacterial groups including potential GABA producers (BAC, LAB, and EREC). Smaller clusters centred on motor-cortex and occipital-cortex GABA+ appeared separate from the peripheral GABA hub in the network layout.

**Figure 4.**
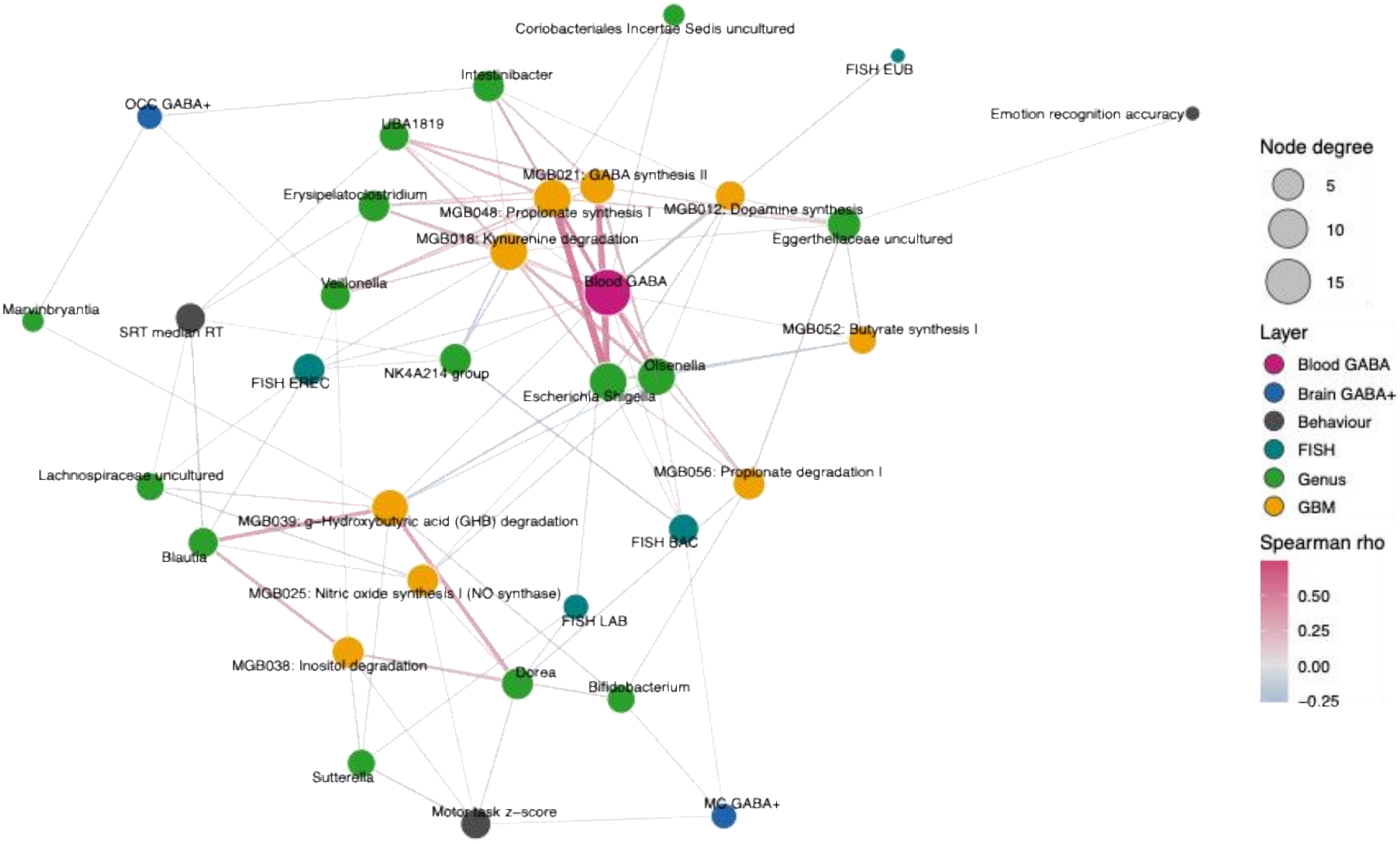
Correlation network linking peripheral (blood) GABA, brain GABA+, behavioural outcomes, and gut microbiota. A GABA-centred correlation network constructed to visualise associations within one step of blood, brain GABA+ or behavioural tasks related to GABA-ergic processing (Spearman correlation |ρ| ≥ 0.15, *p* < 0.05, *q* < 0.10). Nodes represent blood GABA (pink), brain GABA+ (blue), behavioural measures (grey), bacterial genera (green), flow-FISH quantified bacteria groups (teal), and predicted gut-brain modules (GBMs; orange). Node size reflects node degree (number of connections). Edge colour and width represent the strength and direction of the Spearman correlation coefficient (rho), with pink indicating positive and blue indicating negative associations.

#### Sensitivity analyses

Sensitivity analyses supported robustness of the main findings. Additional adjustment for diet-related variables or excluding participants with high autistic traits retained the significant peripheral GABA and brain MRS GABA associations. Sensitivity analyses results are shown in Supplementary Table S8.

## Discussion

This study in 251 healthy adults revealed that gut microbial groups and specific metabolic pathways were associated with both serum GABA and MRS-derived brain GABA. A distinct, but overlapping, pattern of association was found between serum and brain measures, suggesting that peripheral and central GABA may be linked differentially to gut microbiota. Peripheral GABA was associated with quantitative estimates of bacterial groups with recognised potential for GABA production, particularly LAB and BAC (using Flow-FISH). GABA in the motor cortex also showed a positive association with BAC.

The BAC probe in Flow-FISH targets the *Bacteroides*/*Prevotella* group, which includes GABA-producing taxa, such as *Bacteroides fragilis, B. ovatus*, and *B. thetaiotaomicron* (38). These bacteria possess the GAD system and can produce GABA under acidic environments from glutamate. Concentrations of GABA produced by human gut bacteria are within a physiologically bioactive range, suggesting that microbial GABA production may be sufficient to influence local gut signaling(39). Furthermore, the abundance of *Prevotella* CAG:755 have been associated with hippocampal GABA levels in a mouse model of postpartum depression (40). Greater absolute abundance of these bacteria was associated with both peripheral and central GABA. Motor cortex Glx, which reflects the combined glutamate-glutamine pool and a principal precursor for GABA synthesis, was negatively associated with the *Bacteroides/Prevotella* group. This finding provides convergent evidence supporting the observed association between gut microbial composition and motor cortex GABA concentrations.

Both GABA or Glx associations were specific to the motor cortex, indicating regional specificity of these relationships. GABA levels exhibit substantial regional variability across the human brain, with the motor cortex showing the lowest coefficient of variation and the occipital cortex showing the highest (41). Our findings are consistent with experimental evidence in non-human model systems suggesting that gut microorganisms may influence central GABAergic signaling in a region-dependent way. Bravo et al. [5] showed that administration of *Lactobacillus rhamnosus* induced region-specific changes in central GABA receptor expression in rats.

Motor function was related to relative abundance of *Sutterella* as well as the inositol degradation pathway (Figure 2A and 2B). Higher abundance of *Sutterella* has been linked with better psychomotor function in young children as well as in mouse models (42, 43). In contrast, motor network functional connectivity was not associated with any Flow-FISH or 16S-derived microbial measures, suggesting that this GABA-related measure may be less sensitive to microbiota variation. Together, these region- and domain-specific patterns reinforce the notion that the gut microbiota may influence distinct aspects of central nervous system function through multiple, potentially independent pathways, rather than via a single common mechanism.

Peripheral GABA showed a distinct but overlapping microbial association profile compared to central GABA. While the BAC group was positively associated with both peripheral and motor cortex GABA, the LAB group was associated only with peripheral GABA. LAB group includes potentially GABA-producing species, such as *Levilactobacillus brevis* and *Lactiplantibacillus plantarum* (44).These findings suggest that circulating GABA may be linked to additional bacterial communities capable of contributing to GABA-related metabolism.

16S rRNA gene sequencing results identified several genera associated with serum GABA, including *Olsenella* and *Escherichia/Shigella. Escherichia/Shigella* possesses the glutamate decarboxylase system and can produce GABA, and recent evidence has linked higher *Escherichia/Shigella* abundance with increased microbial GABA synthesis (10). This is also supported by our exploratory functional analyses which identified associations with predicted microbial functions related to GABA synthesis and SCFAs metabolism, particularly propionate metabolism. Consistent with these findings, the correlation network showed that peripheral GABA was the most highly connected GABA-related node, with associations involving both microbial taxa and predicted neuroactive microbial functions, particularly propionate and GABA metabolism, while motor GABA+ occupied separate, less connected parts of the network. This pattern suggests that serum GABA is associated with a broader range of microbial and metabolic features, whereas relationships involving regional brain GABA+ appear more distinct. Notably, a recent study in healthy females (45), reported nominal associations between occipital cortex GABA and microbial pathways involved in propionate degradation and p-cresol synthesis. In our study, propionate degradation was positively associated with occipital GABA, while propionate synthesis was associated with peripheral GABA. Together, these findings highlight propionate-related microbial metabolism as a potentially relevant pathway linking the gut microbiota with GABAergic processing.

Given the partially distinct microbiota association patterns observed for peripheral and central GABA, we next examined whether peripheral GABA was associated with MRS-derived brain GABA. To our knowledge, no human studies have directly examined the relationship between circulating GABA concentrations and MRS-derived brain GABA levels. Although there is ongoing debate about whether circulating GABA can cross the blood–brain barrier and influence central nervous system GABA levels, most available evidence comes from animal studies and remains inconclusive (46-48). In our study, no significant association was observed between peripheral and central GABA (Supplementary Table S6). Such a lack of association has also been reported in a rodent model (49). Together, these findings suggest that peripheral GABA is unlikely to represent a reliable proxy for MRS-derived brain GABA. This is consistent with the idea that circulating GABA is influenced by peripheral GABAergic signaling and metabolism (50) and does not necessarily reflect central GABAergic activity. Importantly, lack of association between peripheral and central GABA does not exclude a relationship between the gut microbiota and brain GABA. Gut bacteria may influence central GABAergic function through mechanisms independent of circulating GABA concentrations, including signaling through the vagus nerve (5), modulation of immune and inflammatory pathways (51), endocrine signaling, or the production of other neuroactive metabolites (52). Microbially produced GABA may also act locally on GABA receptors expressed in the enteric nervous system and other intestinal cell types, potentially influencing gut–brain signaling without requiring direct passage across the blood–brain barrier (39).

A major strength of this study is its multi-level characterisation of microbiota-GABA associations in humans. To our knowledge, this is the first study to examine gut microbiota in relation to GABA-related measures across peripheral blood, MRS-derived brain GABA+, motor functional connectivity, and behavioural outcomes linked to GABAergic function. In addition to taxonomic microbiota profiles, we incorporated quantitative bacterial measurements and predicted microbial functional pathways, allowing us to explore whether microbiota associations extended beyond genus-level composition to neuroactive metabolic potential. The robustness of key findings after additional adjustment for dietary factors and after excluding individuals with high autistic traits further supports stability of the observed associations.

However, some minor limitations need to be raised. First, 16S rRNA sequencing provides only a snapshot of gut microbiota taxonomic composition, and specific gene function and regulation related to GABA metabolism could not be directly investigated. An additional limitation is the considerable strain-level variation in GAD system activity. As a result, although taxa associated with central and peripheral GABA levels contain species known to possess the GAD pathway, not all strains within these species are likely to encode or express this system. Accordingly, taxonomic relative abundance may not provide an accurate proxy for in vivo GABA production. However, we used bioinformatic methods to infer metabolic function and identified potential links between the microbiota and several neuroactive compound potentials.

## Conclusion

In conclusion, this study suggests that gut microbiota composition and predicted neuroactive metabolic potential are associated with individual differences in peripheral and brain GABA. Higher cell counts of the *Bacteroides/Prevotella* group were associated with higher MRS-estimated brain GABA, specifically in the motor cortex, as well as peripheral GABA. This highlights the *Bacteroides/Prevotella* group as a promising candidate for future mechanistic studies, particularly given that it contains several known GABA producers. Peripheral GABA showed additional distinct associations with several microbial features, including lactic acid producing bacterial cell counts and predicted bacterial GABA synthesis potential. Together, these findings suggest that microbiota-host GABA relationships may involve broader microbial metabolic and gut-brain signaling pathways, rather than direct microbial GABA production alone. Future longitudinal, metagenomic, metabolomic, and intervention studies are needed to clarify causal direction and underlying mechanisms.

## Acknowledgments

We would like to thank Dr. Carlos Poveda for his assistance with Flow-cytometry analysis and Dr. Nicholas Michael for his assistance at the Chemical Analysis Facility (CAF). We also thank the staff of the Hugh Sinclair Unit, for their assistance with participant recruitment and blood sample collection, and the staff of the Centre for Integrative Neuroscience and Neurodynamics for their help with MRI scans. Finally, we thank all the volunteers who participated in this study. Large Language Model (MS Copilot) was used for grammatical and formatting suggestions.

## Author Contributions

Conceptualization: BC, GG, KAK, AW; Funding Acquisition: BC; Project Administration: CRS, AMM, BC; Methodology: SX, CRS, JK, AMM, VF, CM, SS; Software: CRS, JK, NH, CM; Investigation: SX, CRS, AMM, VF, JN, NK, HR, SS, SD; Data curation: CRS, AMM; Formal Analysis: SX, CRS, AMM, VF; Visualization: SX, CRS, VF; Supervision: BC; Resources: GG, AW, KAK, BC; Writing original draft: SX, AMM; Writing Review and Editing: SX, CRS, AMM, VF, NH, AW, KAK, GG, BC

## Data Availability Statement

All processed data will be shared through the University of Reading Research Data Archive at the point of acceptance. The sequence data obtained by sequencing of the V4-V5 region of the 16S rRNA gene are available from Sequence Read Archive (SRA) of NCBI (https://www.ncbi.nlm.nih.gov/sra) under accession number PRJNA1501397.

