## Supplementary Material for "Gut Microbiota Signatures of Brain and Peripheral gamma-amino butyric acid (GABA) in Healthy Adults"

[a] School of Psychology and Clinical Language Sciences, University of Reading, Reading, RG6 6ES, UK
[b] Department of Food and Nutritional Sciences. School of Chemistry, Food and Pharmacy. University of Reading, Reading, RG6 6UR, UK
[c] Department of Brain & Cognitive Engineering. Korea University, Seoul, South Korea
[d] Brain Research Imaging Centre, Cardiff University, Cardiff, CF24 4HQ, UK

* Prof Bhismadev Chakrabarti
School of Psychology and Clinical Language Sciences
University of Reading
Reading RG6 6ES

**This PDF file includes:**

Supplementary Material and Methods
SI References
Tables S1 to S8

**Supplementary Material and Methods**

*1. Sample collection*

Participants were provided with a faecal sample collection kit containing a sterile collection pot and ice pack. They were instructed to collect a fresh faecal sample at home, preferably on the day of the study visit, and return it within 3 h of collection where possible. Samples were transported with an ice pack, processed upon receipt, and stored at −80°C until analysis. Venous blood samples (9 mL) were collected into serum tubes by a trained phlebotomist during the study visit. Samples were centrifuged at 3,000 × g for 10 min, and the resulting serum was aliquoted and stored at −80°C until analysis. Blood pressure was also measured during the study visit using a standard sphygmomanometer.

*2. Questionnaires assessment*

Anxiety was assessed using the State-Trait Anxiety Inventory (STAI), including state and trait anxiety subscales [1]. Depressive symptoms were assessed using the Centre for Epidemiological Studies Depression Scale [2]. Autistic traits were assessed using the Autism-Spectrum Quotient [3]. Higher scores indicated greater levels of anxiety, depressive symptoms or autistic traits, respectively.

*3. Behavioural assessments*

Participants completed three computer-based behavioural tasks assessing motor performance, emotion recognition, and tactile sensory function. All tasks were completed in a laboratory within the Department of Psychology at University of Reading. Motor performance was assessed using a random motor sequence task, in which participants responded to visual cues by pressing the corresponding keyboard key as quickly and accurately as possible. The motor sequence task included four blocks of 40 trials, and mean reaction time from blocks with accuracy >80% was used as the measure of motor performance. Emotion recognition was assessed using a facial emotion recognition task. Participants viewed faces displaying anger, sadness, happiness, disgust, fear, surprise or neutral expressions and identified the emotion using keyboard responses. The emotion recognition task included one block of 70 trials, and accuracy and mean reaction time for each emotion were extracted. Tactile sensitivity was assessed using the CM4 four-digit tactile stimulator (Cortical Metrics, Carrboro, NC, USA). Vibrotactile stimuli were delivered to the index and middle fingers of the left hand, and participants responded using a mouse with their right hand. The tactile battery included simple reaction time and tactile detection threshold tasks.

Composite scores were derived separately for each behavioural domain (motor, emotion recognition, tactile) using principal component analysis (PCA). Reaction-time variables were reverse-coded so that, for all behavioural variables, higher values indicated better performance before PCA. Variables were then residualised for age, standardised, and entered into PCA. The first principal component was extracted as the task-specific composite score, explaining 78.7%, 32.8%, and 32.5% of variance for the motor, emotion recognition, and tactile domains, respectively. Composite scores were rescaled to range from 0 to 100, with higher scores indicating better overall task performance.

*4. Resting-state functional MRI preprocessing*

Rs-fMRI data were preprocessed with fMRIPrep (version 25.2.3) [4], which performed head-motion correction, susceptibility-distortion correction, co-registration of the functional data to each participant's anatomical image, and spatial normalisation to the MNI152NLin2009cAsym standard space [5] using the transformations distributed via TemplateFlow [6].

*5. MRS data processing*

Spectral modelling was performed using Osprey's built-in linear combination modelling algorithm with the *Separate* fitting approach for edited spectra. The default Osprey basis set was used, with metabolite spectra fitted over the range 0.5–4.0 ppm and water spectra over the range 2.0–7.4 ppm. Baseline modelling employed a spline knot spacing of 0.55 ppm. Macromolecule basis functions were included using Osprey's 1:1 GABA constraint, with the MM3co linewidth fixed at 14Hz. Water-scaled metabolite quantification used an additional unsuppressed water acquisition, while tissue correction was based on segmentation of T1-weighted structural images.

Alpha-corrected concentrations, referenced to the unsuppressed tissue water signal and corrected for voxel grey matter, white matter, and CSF fractions, were calculated as the primary metric, in line with current best-practice recommendations for GABA-edited MRS (Harris et al., 2015; Near et al., 2021). Preliminary correlation analyses revealed that GABA and GABA+ estimates were perfectly correlated ($r = 1$, $p < .001$) across all regions and quantification methods, indicating that the co-edited macromolecule signal included in the GABA+ estimate scaled linearly with the GABA signal in this dataset. Given this redundancy, only the GABA+ estimates were retained for all subsequent analyses.

*6. Post-processing of rsfMRI dataset and functional connectivity measurement (XCP-D)*

Preprocessed BOLD data were post-processed with XCP-D (version 0.13.0) [7]. In brief, XCP-D performed nuisance regression to attenuate motion and physiology-related artefact, temporal filtering, and censoring of high-motion volumes, before extracting mean regional time series and computing functional connectivity. Region-to-region connectivity was estimated as the Pearson correlation between the mean BOLD time series of every pair of parcels, yielding a symmetric connectivity (relationship) matrix per participant. The 1056-region atlas-4S parcellation was used (Schaefer 1000-parcel cortical parcellation augmented with subcortical and cerebellar nodes) [8, 9], in MNI152NLin2009cAsym space.

*7. Motor subnetwork definition and extraction*

The motor subnetwork was defined using the Schaefer 1000-parcel cortical parcellation combined with subcortical and cerebellar nodes (total 1056 parcels). 194 Motor nodes were identified from the atlas parcel labels by their explicit somatomotor-network affiliation, and applied identically across all participants to subset each participant's full 1056 × 1056 connectivity matrix to a 194 × 194 motor-by-motor connectivity submatrix. Within-network motor connectivity was summarised as mean_within_fc_z: each Pearson correlation in the 194 × 194 motor submatrix was Fisher r-to-z transformed, and the mean was taken across all unique off-diagonal edges (upper triangle, excluding self-connections) to yield a single within-motor-network connectivity value per participant. The resulting values for 246 participants formed the basis of the subsequent analyses.

*8. Peripheral GABA measurement*

Serum samples were thawed on ice, and 300 µL of serum was used for sample preparation. Samples, standards and kit-provided low and high controls underwent the derivatisation procedure included in the assay protocol prior to analysis.
A standard curve ranging from 25 to 2,500 ng/mL was prepared according to the manufacturer's recommendations and used for quantification. Absorbance was measured at 450 nm using a microplate reader, and GABA concentrations were calculated by interpolation from the standard curve. The intra-assay coefficient of variation (CV) for the controls was <10%, and the inter-assay CV was ≤15%. To minimise inter-plate batch effects, GABA concentrations were normalised by

dividing each sample value by the mean GABA concentration of the control samples analysed on the same assay plate.

*9. DNA extraction and sequencing*

Briefly, stool samples were mechanically lysed using a TissueLyser LT (Qiagen, Manchester, UK) at 30 Hz for 6 min, followed by DNA extraction and purification according to the manufacturer's protocol.
Extracted DNA samples were sent to Novogene Europe (Cambridge, UK) for 16S rRNA gene sequencing. A 393 bp fragment was amplified using primers 515F, 5′-GTGCCAGCMGCCGCGGTAA-3′, and 907R, 5′-CCGTCAATTCCTTTGAGTTT-3′, together with Phusion® High-Fidelity PCR Master Mix (New England Biolabs, UK).
PCR products were purified, and sequencing libraries were prepared using the NEBNext® Ultra™ DNA Library Prep Kit for Illumina® according to the manufacturer's instructions. Libraries were sequenced on an Illumina NovaSeq 6000 platform using a paired-end 250 bp sequencing strategy, with a target sequencing depth of approximately 30,000 tags per sample. Raw sequencing data were demultiplexed according to sample-specific barcodes by Novogene.

**Table S1.** Partial Spearman correlations adjusted for age, body-fat percentage, and alcohol intake between flow FISH targeted bacterial groups and GABA related variables (p= nominal p-value, q= FDR corrected p-value).

| fish_var | gaba_var | n | partial_rho | p | q |
|---|---|---|---|---|---|
| log_total_EUB | Peripheral GABA | 229 | 0.1990978 | 0.00264136 | 0.01848955 |
| log_total_BIF | Peripheral GABA | 227 | 0.01827355 | 0.78563279 | 0.78563279 |
| log_total_LAB | Peripheral GABA | 225 | 0.17066563 | 0.01085847 | 0.02268992 |
| log_total_BAC | Peripheral GABA | 229 | 0.16506052 | 0.01296567 | 0.02268992 |
| log_total_EREC | Peripheral GABA | 229 | 0.17337658 | 0.00900727 | 0.02268992 |
| log_total_RREC | Peripheral GABA | 229 | 0.09954232 | 0.13573612 | 0.15835881 |
| log_total_CHIS | Peripheral GABA | 220 | 0.12379405 | 0.06874758 | 0.09624662 |
| log_total_EUB | MC GABA+ | 235 | 0.1218925 | 0.06381066 | 0.22333733 |
| log_total_BIF | MC Glx | 232 | -0.0486905 | 0.4634223 | 0.4634223 |
| log_total_LAB | MC Glx | 230 | 0.10612689 | 0.1107923 | 0.25851536 |
| log_total_BAC | MC Glx | 235 | 0.15463239 | 0.01843508 | 0.12904555 |
| log_total_EREC | MC Glx | 235 | 0.08919407 | 0.1757601 | 0.30758017 |
| log_total_RREC | MC Glx | 235 | 0.05268243 | 0.42449026 | 0.4634223 |
| log_total_CHIS | MC Glx | 224 | 0.05214189 | 0.44054456 | 0.4634223 |
| log_total_EUB | MC GABA+ | 236 | 0.01245994 | 0.84995399 | 0.99522682 |
| log_total_BIF | MC GABA+ | 233 | -0.0236757 | 0.72097659 | 0.99522682 |
| log_total_LAB | MC GABA+ | 231 | -0.0515943 | 0.43816747 | 0.99522682 |
| log_total_BAC | MC GABA+ | 236 | 0.00237133 | 0.97128063 | 0.99522682 |
| log_total_EREC | MC Glx | 236 | 0.00039403 | 0.99522682 | 0.99522682 |
| log_total_RREC | MC GABA+ | 236 | -0.1220465 | 0.06289991 | 0.44029937 |
| log_total_CHIS | MC GABA+ | 225 | 0.01093042 | 0.87134918 | 0.99522682 |
| log_total_EUB | OCC GABA+ | 236 | -0.1200558 | 0.06734895 | 0.11135068 |
| log_total_BIF | OCC GABA+ | 233 | -0.0296846 | 0.65427169 | 0.65427169 |
| log_total_LAB | OCC Glx | 231 | -0.1106922 | 0.09544344 | 0.11135068 |
| log_total_BAC | OCC Glx | 236 | -0.1413211 | 0.03105206 | 0.11135068 |
| log_total_EREC | OCC GABA+ | 236 | -0.114593 | 0.080892 | 0.11135068 |
| log_total_RREC | OCC GABA+ | 236 | -0.137429 | 0.03604329 | 0.11135068 |
| log_total_CHIS | OCC GABA+ | 225 | -0.1236514 | 0.06591114 | 0.11135068 |
| log_total_EUB | OCC GABA+ | 236 | -0.0005715 | 0.99307723 | 0.99307723 |
| log_total_BIF | OCC GABA+ | 233 | 0.10872882 | 0.10000076 | 0.7000053 |
| log_total_LAB | OCC Glx | 231 | 0.01971586 | 0.76715713 | 0.89501665 |
| log_total_BAC | OCC Glx | 236 | -0.0254966 | 0.69863913 | 0.89501665 |
| log_total_EREC | OCC Glx | 236 | -0.0719461 | 0.27407758 | 0.89501665 |
| log_total_RREC | OCC Glx | 236 | -0.0457612 | 0.48697865 | 0.89501665 |
| log_total_CHIS | OCC Glx | 225 | 0.03426447 | 0.61159628 | 0.89501665 |
| log_total_EUB | Motor network FC | 235 | -0.0307851 | 0.64086963 | 0.98097382 |
| log_total_BIF | Motor network FC | 232 | -0.0367477 | 0.58010053 | 0.98097382 |
| log_total_LAB | Motor network FC | 230 | -0.0334959 | 0.61565278 | 0.98097382 |
| log_total_BAC | Motor network FC | 235 | 0.07945503 | 0.22797442 | 0.98097382 |
| log_total_EREC | Motor network FC | 235 | 0.00957081 | 0.88471549 | 0.98097382 |
| log_total_RREC | Motor network FC | 235 | 0.0015742 | 0.98097382 | 0.98097382 |
| log_total_CHIS | Motor network FC | 224 | -0.0220786 | 0.74412163 | 0.98097382 |

**Table S2:** Significant (q<0.25) associations between bacterial genera and GABA related variables, adjusted for age, alcohol and body fat percentage.

| feature | metadata | coef | stderr | pval | qval |
|---|---|---|---|---|---|
| Enterobacteriaceae.g__Escherichia.Shigella | Peripheral GABA | 0.323021624 | 0.076149301 | 3.24E-05 | 0.015049289 |
| Atopobiaceae.g__Olsenella | Peripheral GABA | 0.287407809 | 0.091823517 | 0.00198061 | 0.18380061 |
| Coriobacteriales_Incertae_Sedis.g__uncultured | Peripheral GABA | 0.274830339 | 0.090333194 | 0.00262702 | 0.203156194 |
| Lachnospiraceae.g__.Eubacterium._ventriosum_group | Peripheral GABA | -0.301294104 | 0.107488936 | 0.005506432 | 0.21365841 |
| Bifidobacteriaceae.g__Bifidobacterium | MC GABA+ | -0.427821947 | 0.1367548 | 0.001983752 | 0.206413038 |
| Lachnospiraceae.g__Marvinbryantia | OCC GABA+ | 0.277494046 | 0.086397149 | 0.001505794 | 0.150699382 |
| Peptostreptococcaceae.g__Intestinibacter | OCC GABA+ | -0.284884914 | 0.094923113 | 0.002982917 | 0.179591098 |
| Veillonellaceae.g__Veillonella | OCC GABA+ | -0.269389175 | 0.090293086 | 0.003154299 | 0.179591098 |
| Erysipelotrichaceae.g__Turicibacter | OCC GABA+ | -0.293550751 | 0.1071581 | 0.006633076 | 0.240172634 |

**Table S3:** Partial Spearman correlations adjusted for age, body-fat percentage, and alcohol intake between flow FISH targeted bacterial groups and behaviour tasks related to GABA-ergic processing.

| fish_var | cog_var | n | partial_rho | p |
|---|---|---|---|---|
| log_total_EUB | DDT threshold | 226 | 0.101656306 | 0.13016682 |
| log_total_RREC | Emotion-recognition correct RT | 238 | -0.097494986 | 0.1361841 |
| log_total_LAB | Emotion-recognition accuracy | 233 | -0.092204931 | 0.16340554 |
| log_total_EREC | Motor function RT (z) | 235 | 0.085260251 | 0.1956694 |
| log_total_BAC | SRT median RT | 223 | 0.07880968 | 0.24438801 |
| log_total_BIF | Emotion-recognition correct RT | 235 | -0.067348863 | 0.30704284 |
| log_total_LAB | SAD threshold | 218 | 0.069256627 | 0.31211418 |
| log_total_LAB | DDT threshold | 221 | 0.064433277 | 0.34370969 |
| log_total_LAB | Motor function RT (z) | 230 | -0.057280344 | 0.3903645 |
| log_total_EUB | Motor function RT (z) | 235 | 0.05657025 | 0.39106622 |
| log_total_EREC | DDT threshold | 226 | 0.057620313 | 0.39181481 |
| log_total_LAB | Emotion-recognition correct RT | 233 | -0.055121189 | 0.40539067 |
| log_total_EUB | Emotion-recognition correct RT | 238 | -0.054265571 | 0.40764215 |
| log_total_RREC | Emotion-recognition accuracy | 238 | 0.047364192 | 0.46991745 |
| log_total_EREC | Emotion-recognition correct RT | 238 | -0.044563391 | 0.49660641 |
| log_total_BAC | Emotion-recognition correct RT | 238 | 0.041338721 | 0.52830135 |
| log_total_CHIS | SRT median RT | 212 | 0.038759461 | 0.57739906 |
| log_total_EREC | SRT median RT | 223 | -0.037254788 | 0.58257951 |
| log_total_CHIS | Emotion-recognition correct RT | 227 | -0.032735924 | 0.62602026 |
| log_total_EREC | SAD threshold | 223 | -0.031659267 | 0.64048211 |
| log_total_RREC | SRT median RT | 223 | 0.029059044 | 0.66817763 |
| log_total_LAB | SRT median RT | 218 | 0.028279655 | 0.68010025 |
| log_total_RREC | Motor function RT (z) | 235 | 0.021849764 | 0.74060852 |
| log_total_BIF | SRT median RT | 220 | 0.022046909 | 0.74674468 |
| log_total_BAC | DDT threshold | 226 | 0.01968613 | 0.77001687 |
| log_total_EUB | SAD threshold | 223 | 0.014068497 | 0.83562637 |
| log_total_CHIS | SAD threshold | 213 | 0.013702253 | 0.84352345 |
| log_total_BAC | Motor function RT (z) | 235 | 0.011661978 | 0.85976248 |
| log_total_BIF | Motor function RT (z) | 232 | 0.011309695 | 0.86484057 |
| log_total_BAC | SAD threshold | 223 | -0.010168044 | 0.88079525 |
| log_total_CHIS | Emotion-recognition accuracy | 227 | 0.009677634 | 0.88547316 |
| log_total_EUB | SRT median RT | 223 | 0.009142561 | 0.89274123 |
| log_total_CHIS | Motor function RT (z) | 224 | -0.00808428 | 0.90487775 |
| log_total_BAC | Emotion-recognition accuracy | 238 | -0.007314173 | 0.91119784 |
| log_total_RREC | DDT threshold | 226 | -0.006974797 | 0.91750913 |
| log_total_BIF | SAD threshold | 220 | -0.006478178 | 0.92441067 |
| log_total_BIF | DDT threshold | 223 | 0.006225853 | 0.92684173 |
| log_total_CHIS | DDT threshold | 215 | -0.006118623 | 0.92942952 |
| log_total_EREC | Emotion-recognition accuracy | 238 | -0.005392859 | 0.93446323 |
| log_total_RREC | SAD threshold | 223 | 0.005439589 | 0.93605981 |
| log_total_BIF | Emotion-recognition accuracy | 235 | 0.005157442 | 0.93772304 |
| log_total_EUB | Emotion-recognition accuracy | 238 | 0.004826796 | 0.94132925 |

**Table S4:** Significant (q<0.25) associations between bacterial genera and behaviour tasks related to GABA-ergic processing, adjusted for age, alcohol and body fat percentage.

| feature | metadata | coef | stderr | pval | qval |
|---|---|---|---|---|---|
| Eggerthellaceae.g__uncultured | Emotion accuracy | -0.3162138 | 0.11526519 | 0.00655236 | 0.24281779 |
| Lachnospiraceae.g__Lachnoclostridium | Emotion accuracy | 0.17056631 | 0.06347199 | 0.00772095 | 0.24281779 |
| Erysipelatoclostridiaceae.g__Erysipelatoclostridium | SRT median RT | -0.2472075 | 0.07357428 | 0.00091949 | 0.09111509 |
| Lachnospiraceae.g__uncultured | SRT median RT | -0.2260426 | 0.07846324 | 0.00435986 | 0.18986499 |
| Oscillospiraceae.g__NK4A214_group | SRT median RT | 0.34862272 | 0.12145351 | 0.00450111 | 0.18986499 |
| Ruminococcaceae.g__uncultured | SRT median RT | 0.25758626 | 0.09345192 | 0.00633776 | 0.20573207 |
| Lachnospiraceae.g__Blautia | SRT median RT | -0.1476099 | 0.0542401 | 0.00702306 | 0.20573207 |
| Ruminococcaceae.g__UBA1819 | SRT median RT | -0.1467887 | 0.05692045 | 0.01056812 | 0.24518045 |
| Sutterellaceae.g__Sutterella | Motor task z-score | -0.5432422 | 0.1457481 | 0.00024347 | 0.03823452 |
| Lachnospiraceae.g__Dorea | Motor task z-score | 0.16989134 | 0.05634018 | 0.00285272 | 0.17923447 |
| Sutterellaceae.g__Sutterella | Motor composite | 0.53192182 | 0.13916336 | 0.00016936 | 0.03833915 |

**Table S5:** Nominally significant ($p<0.05$) associations between predicted gut brain modules and GABA related variables, adjusted for age, alcohol and body fat percentage.

| feature | metadata | coef | stderr | pval | qval | type |
|---|---|---|---|---|---|---|
| MGB038: Inositol degradation | Motor function RT (z) | 0.139521581 | 0.04964778 | 0.0053753 | 0.22629183 | Motor |
| MGB025: Nitric oxide synthesis I (NO synthase) | Motor function RT (z) | -0.293004806 | 0.11667235 | 0.01271109 | 0.2440977 | Motor |
| MGB027: Nitric oxide degradation I (NO dioxygenase) | Motor function RT (z) | -0.171334178 | 0.0706018 | 0.01600034 | 0.2440977 | Motor |
| MGB009: Histamine synthesis | Motor function RT (z) | -0.215545085 | 0.09892039 | 0.03034476 | 0.30701761 | Motor |
| MGB032: Quinolinic acid synthesis | Motor function RT (z) | 0.051472625 | 0.02430115 | 0.03523386 | 0.33647971 | Motor |
| MGB039: g-Hydroxybutyric acid (GHB) degradation | Motor function RT (z) | 0.082989544 | 0.04134782 | 0.04590337 | 0.34087457 | Motor |
| MGB012: Dopamine synthesis | Emotion-recognition correct RT | 0.242387606 | 0.11113756 | 0.03018205 | 0.37849105 | Emotion recognition |
| MGB025: Nitric oxide synthesis I (NO synthase) | SAD threshold | -0.276679729 | 0.11362397 | 0.01568996 | 0.38401252 | Tactile |
| MGB024: DOPAC synthesis | SAD threshold | 0.570112107 | 0.26299858 | 0.03125683 | 0.38401252 | Tactile |
| MGB055: Propionate synthesis III | SAD threshold | -0.095063368 | 0.04562701 | 0.03836698 | 0.42712173 | Tactile |
| MGB028: Nitric oxide degradation II (NO reductase) | DDT threshold | -0.153550835 | 0.06975686 | 0.02874994 | 0.44954456 | Tactile |
| MGB025: Nitric oxide synthesis I (NO synthase) | DDT threshold | -0.240577015 | 0.11369116 | 0.03545384 | 0.47562415 | Tactile |
| MGB047: Acetate degradation | DDT threshold | -0.084801393 | 0.04294769 | 0.04956134 | 0.47562415 | Tactile |
| MGB048: Propionate synthesis I | Peripheral GABA | 0.191206543 | 0.05320119 | 0.00040037 | 0.068864 | Peripheral GABA |
| MGB018: Kynurenine degradation | Peripheral GABA | 0.184739474 | 0.06851382 | 0.00754223 | 0.14287918 | Peripheral GABA |
| MGB056: Propionate degradation I | Peripheral GABA | 0.251247688 | 0.08796623 | 0.00469084 | 0.14287918 | Peripheral GABA |
| MGB021: GABA synthesis II | Peripheral GABA | 0.389004159 | 0.1427146 | 0.00692278 | 0.14287918 | Peripheral GABA |
| MGB012: Dopamine synthesis | Peripheral GABA | -0.257602999 | 0.10927223 | 0.01926282 | 0.21858606 | Peripheral GABA |
| MGB015: p-Cresol synthesis | Peripheral GABA | -0.0512163 | 0.02447447 | 0.03750702 | 0.22245544 | Peripheral GABA |
| MGB039: g-Hydroxybutyric acid (GHB) degradation | Peripheral GABA | -0.093247304 | 0.04185409 | 0.02688035 | 0.22245544 | Peripheral GABA |
| MGB034: Isovaleric acid synthesis I (KADH pathway) | Peripheral GABA | -0.149457518 | 0.06877729 | 0.03082426 | 0.22245544 | Peripheral GABA |
| MGB052: Butyrate synthesis I | Peripheral GABA | -0.073563528 | 0.0331982 | 0.02770533 | 0.22245544 | Peripheral GABA |

**Table S6:** Spearman correlations between peripheral GABA and brain GABA.

| Peripheral | Brain | n | rho | p | q |
|---|---|---|---|---|---|
| GABA | MC GABA+ | 236 | -0.0892645 | 0.17170159 | 0.33461222 |
| Gln | MC GABA+ | 238 | -0.1401063 | 0.03071511 | 0.15461017 |
| Glu | MC GABA+ | 234 | -0.0251942 | 0.70142702 | 0.80933887 |
| GABA | OCC GABA+ | 236 | -0.0599904 | 0.35886871 | 0.53830307 |
| Gln | OCC GABA+ | 238 | 0.14084515 | 0.02983658 | 0.15461017 |
| Glu | OCC GABA+ | 234 | -0.083119 | 0.20520505 | 0.34200841 |
| GABA | MC Glx | 237 | -0.1192107 | 0.06694233 | 0.25103375 |
| Gln | MC Glx | 239 | 0.10765107 | 0.09684439 | 0.29053316 |
| Glu | MC Glx | 235 | -0.0321827 | 0.62353427 | 0.77941784 |
| GABA | OCC Glx | 236 | -0.0089265 | 0.89150026 | 0.89150026 |
| Gln | OCC Glx | 238 | 0.08750993 | 0.17845985 | 0.33461222 |
| Glu | OCC Glx | 234 | 0.14112666 | 0.03092203 | 0.15461017 |
| GABA | Motor network FC | 235 | -0.0354504 | 0.58870312 | 0.77941784 |
| Gln | Motor network FC | 237 | -0.0157842 | 0.80899275 | 0.86677795 |
| Glu | Motor network FC | 233 | 0.09300341 | 0.15704817 | 0.33461222 |

**Table S7:** Spearman correlations for peripheral (blood) GABA, brain GABA and gut microbiota features correlation network

| from_label | to_label | rho | p | q | abs |
|---|---|---|---|---|---|
| Blood GABA | FISH BAC | 0.15996252 | 0.01309451 | 0.04588674 | 0.15996252 |
| Blood GABA | FISH EUB | 0.21069694 | 0.00102343 | 0.00578768 | 0.21069694 |
| Blood GABA | FISH LAB | 0.18669644 | 0.00399944 | 0.01777619 | 0.18669644 |
| Blood GABA | FISH EREC | 0.18855591 | 0.00336579 | 0.0155053 | 0.18855591 |
| Blood GABA | Erysipelatoclostridium | 0.2098675 | 0.00107251 | 0.00598272 | 0.2098675 |
| Blood GABA | NK4A214 group | -0.1654722 | 0.01023384 | 0.03746318 | 0.16547216 |
| Blood GABA | UBA1819 | 0.17199138 | 0.0075739 | 0.02929527 | 0.17199138 |
| Blood GABA | Intestinibacter | 0.20224409 | 0.00163604 | 0.00854494 | 0.20224409 |
| Blood GABA | Escherichia Shigella | 0.32264904 | 3.23E-07 | 4.56E-06 | 0.32264904 |
| Blood GABA | Olsenella | 0.34656093 | 3.53E-08 | 6.59E-07 | 0.34656093 |
| Blood GABA | Coriobacteriales Incertae Sedis uncultured | 0.18866572 | 0.00334699 | 0.0155053 | 0.18866572 |
| Blood GABA | MGB048: Propionate synthesis I | 0.33723611 | 8.56E-08 | 1.43E-06 | 0.33723611 |
| Blood GABA | MGB056: Propionate degradation I | 0.27351317 | 1.73E-05 | 0.00017941 | 0.27351317 |
| Blood GABA | MGB021: GABA synthesis II | 0.28407803 | 7.81E-06 | 8.89E-05 | 0.28407803 |
| Blood GABA | MGB018: Kynurenine degradation | 0.23538662 | 0.00023379 | 0.00172713 | 0.23538662 |
| Blood GABA | MGB012: Dopamine synthesis | -0.2595993 | 4.69E-05 | 0.00043675 | 0.25959929 |
| Blood GABA | MGB039: g-Hydroxybutyric acid (GHB) degradation | -0.1772402 | 0.00589997 | 0.02403747 | 0.17724017 |
| Blood GABA | MGB052: Butyrate synthesis I | -0.1580597 | 0.01423433 | 0.04850589 | 0.1580597 |
| MC GABA+ | Motor task z-score | -0.1642153 | 0.01018692 | 0.03745861 | 0.16421529 |
| MC GABA+ | FISH BAC | 0.15629062 | 0.0141302 | 0.04850589 | 0.15629062 |
| MC GABA+ | Bifidobacterium | -0.1679429 | 0.00817263 | 0.03117005 | 0.16794293 |
| OCC GABA+ | Marvinbryantia | 0.19595014 | 0.0019746 | 0.01011982 | 0.19595014 |
| OCC GABA+ | Veillonella | -0.1521324 | 0.01672196 | 0.05573987 | 0.15213243 |
| OCC GABA+ | Intestinibacter | -0.1905279 | 0.00264028 | 0.0128871 | 0.19052788 |
| Emotion recognition accuracy | Eggerthellaceae uncultured | -0.1543839 | 0.01474771 | 0.04935968 | 0.15438388 |
| SRT median RT | Erysipelatoclostridium | -0.1748144 | 0.00747956 | 0.02920589 | 0.17481441 |
| SRT median RT | NK4A214 group | 0.15384196 | 0.01879001 | 0.06114209 | 0.15384196 |
| SRT median RT | Lachnospiraceae uncultured | -0.1613283 | 0.01368231 | 0.04754022 | 0.16132833 |
| SRT median RT | Blautia | -0.224526 | 0.00055439 | 0.00357951 | 0.22452604 |
| SRT median RT | UBA1819 | -0.1709598 | 0.00892715 | 0.03342588 | 0.17095975 |
| Motor task z-score | Sutterella | -0.220721 | 0.00047494 | 0.00319224 | 0.22072104 |
| Motor task z-score | Dorea | 0.2122488 | 0.00078747 | 0.00461231 | 0.2122488 |
| Motor task z-score | MGB038: Inositol degradation | 0.17502477 | 0.00581386 | 0.02383684 | 0.17502477 |
| Motor task z-score | MGB025: Nitric oxide synthesis I (NO synthase) | -0.15909 | 0.01229444 | 0.04345449 | 0.15909004 |
| FISH BAC | NK4A214 group | -0.2458688 | 8.55E-05 | 0.00073007 | 0.24586877 |
| FISH BAC | Escherichia Shigella | 0.15445226 | 0.01450319 | 0.04874023 | 0.15445226 |
| FISH BAC | Olsenella | 0.17264955 | 0.00620533 | 0.02494299 | 0.17264955 |
| FISH LAB | Sutterella | -0.1753576 | 0.00592142 | 0.02403747 | 0.17535761 |
| FISH LAB | Dorea | -0.1842422 | 0.00380473 | 0.01723689 | 0.18424222 |
| FISH EREC | Erysipelatoclostridium | 0.16287582 | 0.00989047 | 0.0368082 | 0.16287582 |
| FISH EREC | NK4A214 group | -0.1671659 | 0.00808398 | 0.03097602 | 0.16716594 |
| FISH EREC | Lachnospiraceae uncultured | 0.15447761 | 0.01448687 | 0.04874023 | 0.15447761 |
| FISH EREC | Blautia | 0.1849124 | 0.00334175 | 0.0155053 | 0.1849124 |
| FISH EREC | MGB018: Kynurenine degradation | 0.18264215 | 0.00375829 | 0.01712112 | 0.18264215 |
| Eggerthellaceae uncultured | MGB048: Propionate synthesis I | 0.24878315 | 6.76E-05 | 0.00058339 | 0.24878315 |
| Eggerthellaceae uncultured | MGB056: Propionate degradation I | 0.23034187 | 0.00023259 | 0.00172713 | 0.23034187 |
| Eggerthellaceae uncultured | MGB021: GABA synthesis II | 0.20797344 | 0.00091712 | 0.00525899 | 0.20797344 |
| Eggerthellaceae uncultured | MGB018: Kynurenine degradation | 0.17467881 | 0.00552041 | 0.02286231 | 0.17467881 |
| Eggerthellaceae uncultured | MGB012: Dopamine synthesis | -0.1744216 | 0.00559127 | 0.02303942 | 0.17442155 |
| Eggerthellaceae uncultured | MGB052: Butyrate synthesis I | -0.2098729 | 0.00082059 | 0.00477222 | 0.20987289 |
| Sutterella | MGB038: Inositol degradation | -0.2031577 | 0.00121059 | 0.00670733 | 0.20315766 |
| Sutterella | MGB039: g-Hydroxybutyric acid (GHB) degradation | -0.1779131 | 0.00469578 | 0.02026601 | 0.17791311 |
| Dorea | MGB056: Propionate degradation I | -0.1949383 | 0.00191693 | 0.00988606 | 0.19493834 |
| Dorea | MGB038: Inositol degradation | 0.28377158 | 4.93E-06 | 5.86E-05 | 0.28377158 |
| Dorea | MGB025: Nitric oxide synthesis I (NO synthase) | -0.1621506 | 0.01007695 | 0.03722117 | 0.16215064 |
| Dorea | MGB039: g-Hydroxybutyric acid (GHB) degradation | 0.46485525 | 7.35E-15 | 3.01E-13 | 0.46485525 |
| Erysipelatoclostridium | MGB048: Propionate synthesis I | 0.24904041 | 6.64E-05 | 0.00058157 | 0.24904041 |
| Erysipelatoclostridium | MGB021: GABA synthesis II | 0.2239484 | 0.00034913 | 0.00244687 | 0.2239484 |
| Erysipelatoclostridium | MGB018: Kynurenine degradation | 0.32699298 | 1.15E-07 | 1.85E-06 | 0.32699298 |
| NK4A214 group | MGB048: Propionate synthesis I | -0.2176854 | 0.00051407 | 0.00342713 | 0.21768545 |
| NK4A214 group | MGB018: Kynurenine degradation | -0.2568833 | 3.81E-05 | 0.00036739 | 0.25688332 |
| Lachnospiraceae uncultured | MGB025: Nitric oxide synthesis I (NO synthase) | -0.2124561 | 0.00070431 | 0.00427804 | 0.21245608 |
| Lachnospiraceae uncultured | MGB039: g-Hydroxybutyric acid (GHB) degradation | 0.20586075 | 0.0010367 | 0.00582254 | 0.20586075 |
| Blautia | MGB038: Inositol degradation | 0.40269171 | 3.34E-11 | 9.12E-10 | 0.40269171 |
| Blautia | MGB025: Nitric oxide synthesis I (NO synthase) | -0.1771611 | 0.00487695 | 0.0206321 | 0.17716107 |
| Blautia | MGB039: g-Hydroxybutyric acid (GHB) degradation | 0.47198862 | 2.49E-15 | 1.14E-13 | 0.47198862 |
| UBA1819 | MGB048: Propionate synthesis I | 0.35231417 | 9.53E-09 | 1.95E-07 | 0.35231417 |
| UBA1819 | MGB021: GABA synthesis II | 0.31075242 | 5.07E-07 | 6.82E-06 | 0.31075242 |
| UBA1819 | MGB018: Kynurenine degradation | 0.30329425 | 9.73E-07 | 1.25E-05 | 0.30329425 |
| Bifidobacterium | MGB056: Propionate degradation I | -0.1771102 | 0.00488943 | 0.0206321 | 0.17711023 |
| Bifidobacterium | MGB038: Inositol degradation | 0.23484045 | 0.00017359 | 0.00138196 | 0.23484045 |
| Bifidobacterium | MGB039: g-Hydroxybutyric acid (GHB) degradation | 0.21075545 | 0.00077901 | 0.00459558 | 0.21075545 |
| Marvinbryantia | MGB039: g-Hydroxybutyric acid (GHB) degradation | 0.17812939 | 0.00464481 | 0.0201521 | 0.17812939 |
| Veillonella | MGB048: Propionate synthesis I | 0.27717625 | 8.31E-06 | 9.21E-05 | 0.27717625 |
| Veillonella | MGB038: Inositol degradation | -0.1550605 | 0.01392273 | 0.04817148 | 0.15506052 |
| Veillonella | MGB021: GABA synthesis II | 0.23234529 | 0.00020432 | 0.00158057 | 0.23234529 |
| Veillonella | MGB018: Kynurenine degradation | 0.24926048 | 6.54E-05 | 0.00058157 | 0.24926048 |
| Intestinibacter | MGB048: Propionate synthesis I | 0.34612332 | 1.79E-08 | 3.41E-07 | 0.34612332 |
| Intestinibacter | MGB021: GABA synthesis II | 0.26377158 | 2.30E-05 | 0.00023031 | 0.26377158 |
| Intestinibacter | MGB018: Kynurenine degradation | 0.16763423 | 0.00778099 | 0.029955 | 0.16763423 |
| Intestinibacter | MGB012: Dopamine synthesis | -0.1590415 | 0.01162938 | 0.04162261 | 0.15904155 |
| Escherichia Shigella | MGB048: Propionate synthesis I | 0.7461347 | 6.93E-46 | 1.90E-43 | 0.7461347 |
| Escherichia Shigella | MGB056: Propionate degradation I | 0.21439803 | 0.00062713 | 0.00392558 | 0.21439803 |
| Escherichia Shigella | MGB021: GABA synthesis II | 0.62456384 | 1.48E-28 | 3.04E-26 | 0.62456384 |
| Escherichia Shigella | MGB018: Kynurenine degradation | 0.30524682 | 8.22E-07 | 1.07E-05 | 0.30524682 |
| Escherichia Shigella | MGB025: Nitric oxide synthesis I (NO synthase) | -0.1505316 | 0.01700593 | 0.05645693 | 0.15053159 |
| Escherichia Shigella | MGB012: Dopamine synthesis | -0.2117063 | 0.00073639 | 0.00443998 | 0.21170632 |
| Escherichia Shigella | MGB039: g-Hydroxybutyric acid (GHB) degradation | -0.2633891 | 2.37E-05 | 0.00023406 | 0.26338911 |
| Escherichia Shigella | MGB052: Butyrate synthesis I | -0.2491709 | 6.58E-05 | 0.00058157 | 0.24917094 |
| Olsenella | MGB048: Propionate synthesis I | 0.47589072 | 1.37E-15 | 6.59E-14 | 0.47589072 |
| Olsenella | MGB056: Propionate degradation I | 0.24417606 | 9.29E-05 | 0.00078512 | 0.24417606 |
| Olsenella | MGB021: GABA synthesis II | 0.34889395 | 1.35E-08 | 2.64E-07 | 0.34889395 |
| Olsenella | MGB018: Kynurenine degradation | 0.41188086 | 1.07E-11 | 3.25E-10 | 0.41188086 |
| Olsenella | MGB025: Nitric oxide synthesis I (NO synthase) | -0.1921078 | 0.00223646 | 0.0111823 | 0.19210776 |
| Olsenella | MGB012: Dopamine synthesis | -0.1770412 | 0.00490641 | 0.0206321 | 0.17704117 |
| Olsenella | MGB039: g-Hydroxybutyric acid (GHB) degradation | -0.182524 | 0.00371138 | 0.01700186 | 0.182524 |
| Olsenella | MGB052: Butyrate synthesis I | -0.1923202 | 0.00221089 | 0.01112225 | 0.19232024 |
| Coriobacteriales Incertae Sedis uncultured | MGB048: Propionate synthesis I | 0.17669588 | 0.00499214 | 0.0208855 | 0.17669588 |

**Table S8:** Sensitivity analysis for GBMs, gut microbiota genera that related to peripheral GABA or brain GABA+.

| **Models: additionally adjusted for enery and fibre intake** | | | | | | |
|---|---|---|---|---|---|---|
| **feature** | **metadata** | **coef** | **stderr** | **pval** | **qval** | **Significance status** |
| Enterobacteriaceae.g__Escherichia.Shigella | Peripheral GABA | 0.31985207 | 0.07687513 | 4.537608204 | 0.03158175 | Retained |
| Atopobiaceae.g__Olsenella | Peripheral GABA | 0.29383204 | 0.09236654 | 0.0016766 | 0.10608301 | Retained |
| Coriobacteriales_Incertae_Sedis.g__uncultured | Peripheral GABA | 0.25816791 | 0.09061915 | 0.00479894 | 0.19220028 | Retained |
| Lachnospiraceae.g__.Eubacterium._ventriosum_group | Peripheral GABA | -0.2989406 | 0.10853874 | 0.00637076 | 0.20245274 | Retained |
| Bifidobacteriaceae.g__Bifidobacterium | MC GABA+ | -0.4511179 | 0.13837951 | 0.00128369 | 0.1061895 | Retained |
| Lachnospiraceae.g__Marvinbryantia | OCC GABA+ | 0.26708096 | 0.0878791 | 0.00264656 | 0.12714073 | Retained |
| Peptostreptococcaceae.g__Intestinibacter | OCC GABA+ | -0.2858859 | 0.09630085 | 0.00330735 | 0.13672272 | Retained |
| Veillonellaceae.g__Veillonella | OCC GABA+ | -0.3067284 | 0.09069735 | 0.00084588 | 0.08410484 | Retained |
| Erysipelotrichaceae.g__Turicibacter | OCC GABA+ | -0.3056631 | 0.10883627 | 0.00540568 | 0.16677118 | Retained |
| MGB048: Propionate synthesis I | Peripheral GABA | 0.18369788 | 0.05209644 | 0.00051221 | 0.13215082 | Retained |
| MGB018: Kynurenine degradation | Peripheral GABA | 0.1809172 | 0.06699225 | 0.00745581 | 0.15226444 | Retained |
| MGB021: GABA synthesis II | Peripheral GABA | 0.35855585 | 0.13935168 | 0.01073276 | 0.15948847 | Retained |
| MGB056: Propionate degradation I | Peripheral GABA | 0.25124878 | 0.08574013 | 0.00373952 | 0.14368827 | Retained |
| MGB012: Dopamine synthesis | Peripheral GABA | -0.2347651 | 0.10564391 | 0.02727665 | 0.23844091 | Retained |
| MGB015: p-Cresol synthesis | Peripheral GABA | -0.0490986 | 0.02395374 | 0.04156674 | 0.23844091 | Retained |
| MGB034: Isovaleric acid synthesis I (KADH pathway) | Peripheral GABA | -0.1464246 | 0.06716903 | 0.03031323 | 0.23844091 | Retained |
| MGB039: g-Hydroxybutyric acid (GHB) degradation | Peripheral GABA | -0.0869111 | 0.04093372 | 0.03484276 | 0.23844091 | Retained |
| MGB052: Butyrate synthesis I | Peripheral GABA | -0.0801186 | 0.03189807 | 0.01272651 | 0.17281255 | Retained |
| **Models: excluding individuals with high potential of autistic traits, adjusted for age, body fat and alcohol intake** | | | | | | |
| **feature** | **metadata** | **coef** | **stderr** | **pval** | **qval** | **Significance status** |
| Enterobacteriaceae.g__Escherichia.Shigella | Peripheral GABA | 0.34080844 | 0.07879499 | 2.363838667 | 0.01096821 | Retained |
| Atopobiaceae.g__Olsenella | Peripheral GABA | 0.28920997 | 0.09815861 | 0.00358208 | 0.23182803 | Retained |
| Coriobacteriales_Incertae_Sedis.g__uncultured | Peripheral GABA | 0.28835822 | 0.09433446 | 0.00253085 | 0.23182803 | Retained |
| Lachnospiraceae.g__.Eubacterium._ventriosum_group | Peripheral GABA | -0.309794 | 0.10969923 | 0.00520348 | 0.23182803 | Retained |
| Bifidobacteriaceae.g__Bifidobacterium | MC GABA+ | -0.5069665 | 0.14072473 | 0.00039108 | 0.13406158 | Retained |
| Lachnospiraceae.g__Marvinbryantia | OCC GABA+ | 0.29279685 | 0.0884069 | 0.00108529 | 0.141819 | Retained |
| Peptostreptococcaceae.g__Intestinibacter | OCC GABA+ | -0.2916851 | 0.09833634 | 0.00335267 | 0.20430884 | Retained |
| Veillonellaceae.g__Veillonella | OCC GABA+ | -0.2789252 | 0.09467204 | 0.00356788 | 0.20430884 | Retained |
| Erysipelotrichaceae.g__Turicibacter | OCC GABA+ | -0.3601223 | 0.10990215 | 0.00122258 | 0.141819 | Retained |
| MGB048: Propionate synthesis I | Peripheral GABA | 0.19793567 | 0.05310169 | 0.00024927 | 0.04287421 | Retained |
| MGB018: Kynurenine degradation | Peripheral GABA | 0.18016859 | 0.06601434 | 0.00689191 | 0.15061308 | Retained |
| MGB021: GABA synthesis II | Peripheral GABA | 0.39148907 | 0.14498681 | 0.00750104 | 0.15061308 | Retained |
| MGB056: Propionate degradation I | Peripheral GABA | 0.22255656 | 0.08780956 | 0.01199557 | 0.16133367 | Retained |
| MGB012: Dopamine synthesis | Peripheral GABA | -0.239549 | 0.11081038 | 0.03177599 | 0.18002815 | Retained |
| MGB015: p-Cresol synthesis | Peripheral GABA | -0.0570478 | 0.02421568 | 0.01941201 | 0.18002815 | Retained |
| MGB034: Isovaleric acid synthesis I (KADH pathway) | Peripheral GABA | -0.1557695 | 0.07054543 | 0.02833158 | 0.18002815 | Retained |
| MGB039: g-Hydroxybutyric acid (GHB) degradation | Peripheral GABA | -0.0933215 | 0.04216234 | 0.02795785 | 0.18002815 | Retained |
| MGB052: Butyrate synthesis I | Peripheral GABA | -0.0722918 | 0.03341017 | 0.03162201 | 0.18002815 | Retained |
| **Models: excluding individuals with high potential of autistic traits, adjusted for age, body fat, alcohol, energy and fibre intake** | | | | | | |
| **feature** | **metadata** | **coef** | **stderr** | **pval** | **qval** | **Significance status** |
| Enterobacteriaceae.g__Escherichia.Shigella | Peripheral GABA | 0.33912184 | 0.07967015 | 3.152363504 | 0.02194045 | Retained |
| Atopobiaceae.g__Olsenella | Peripheral GABA | 0.29373375 | 0.09892637 | 0.00333965 | 0.25826618 | Not significant |
| Coriobacteriales_Incertae_Sedis.g__uncultured | Peripheral GABA | 0.27137317 | 0.09490234 | 0.00467967 | 0.29609538 | Not significant |
| Lachnospiraceae.g__.Eubacterium._ventriosum_group | Peripheral GABA | -0.3061757 | 0.11091314 | 0.00629203 | 0.31519559 | Not significant |
| Bifidobacteriaceae.g__Bifidobacterium | MC GABA+ | -0.5290302 | 0.14252318 | 0.00026235 | 0.12525834 | Retained |
| Lachnospiraceae.g__Marvinbryantia | OCC GABA+ | 0.28836304 | 0.08962306 | 0.00149282 | 0.20780017 | Retained |
| Peptostreptococcaceae.g__Intestinibacter | OCC GABA+ | -0.2892604 | 0.09928976 | 0.0039541 | 0.24826994 | Retained |
| Veillonellaceae.g__Veillonella | OCC GABA+ | -0.3069555 | 0.09496615 | 0.00142125 | 0.20780017 | Retained |
| Erysipelotrichaceae.g__Turicibacter | OCC GABA+ | -0.3607751 | 0.1108156 | 0.00131436 | 0.20780017 | Retained |
| MGB048: Propionate synthesis I | Peripheral GABA | 0.19637594 | 0.05358257 | 0.00031475 | 0.08120433 | Retained |
| MGB018: Kynurenine degradation | Peripheral GABA | 0.18699298 | 0.06649893 | 0.00539976 | 0.18260407 | Retained |
| MGB021: GABA synthesis II | Peripheral GABA | 0.37636468 | 0.14649479 | 0.01090212 | 0.18747482 | Retained |
| MGB056: Propionate degradation I | Peripheral GABA | 0.23148086 | 0.08870173 | 0.00972851 | 0.18747482 | Retained |
| MGB012: Dopamine synthesis | Peripheral GABA | -0.2195373 | 0.11103478 | 0.04935346 | 0.31017443 | Not significant |
| MGB015: p-Cresol synthesis | Peripheral GABA | -0.0577925 | 0.02448996 | 0.0192173 | 0.209539 | Retained |
| MGB034: Isovaleric acid synthesis I (KADH pathway) | Peripheral GABA | -0.1571225 | 0.07113943 | 0.02830103 | 0.22817709 | Retained |
| MGB039: g-Hydroxybutyric acid (GHB) degradation | Peripheral GABA | -0.0891095 | 0.04259866 | 0.03767848 | 0.27002912 | Not significant |
| MGB052: Butyrate synthesis I | Peripheral GABA | -0.0821672 | 0.03312946 | 0.0139331 | 0.18919688 | Retained |
